# Dispersal syndromes can impact ecosystem functioning in spatially structured freshwater populations

**DOI:** 10.1101/485706

**Authors:** Chelsea J. Little, Emanuel A. Fronhofer, Florian Altermatt

**Author notes:** Corresponding author: Department of Aquatic Ecology, Eawag: Swiss Federal Institute of Aquatic Science and Technology, Überlandstrasse 133, 8600 Dübendorf, Switzerland.

## Abstract

Dispersal can strongly influence ecological and evolutionary dynamics. Besides the direct contribution of dispersal to population dynamics, dispersers often differ in their phenotypic attributes from non-dispersers, which leads to dispersal syndromes. The consequences of such dispersal syndromes have been widely explored at the population and community level, however, to date, ecosystem-level effects remain unclear. Here, we examine whether dispersing and resident individuals of two different aquatic keystone invertebrate species have different contributions to detrital processing, a key function in freshwater ecosystems. Using experimental two-patch systems, we found no difference in leaf consumption rates with dispersal status of the common native species *Gammarus fossarum*. In *Dikerogammarus villosus*, however, a Ponto-Caspian species now expanding throughout Europe, dispersers consumed leaf litter at roughly three times the rate of non-dispersers. Furthermore, this put the contribution of dispersing *D. villosus* to leaf litter processing on par with native *G. fossarum,* after adjusting for differences in organismal size. Given that leaf litter decomposition is a key function in aquatic ecosystems, and the rapid species turnover in freshwater habitats with range expansions of non-native species, this finding suggests that dispersal syndromes may have important consequences for ecosystem functioning.

## Introduction

Dispersal, the movement from a natal site to another site or habitat patch with potential consequences for gene flow, is an essential process in ecology and evolution [1,2]. Dispersal connects local populations and allows for colonization of new patches, and thus governs the spatial distribution of biodiversity. Although it is often treated as a stochastic event, dispersal between patches is neither neutral with respect to species [3] nor to individuals within species [4]. Within species, individuals may disperse depending on their own phenotype (dispersal syndrome) [5–7]. Across the animal kingdom, dispersing and non-dispersing individuals have identifiable differences in a broad range of phenotypic characteristics [2,4,8,9].

To date, the consequences of dispersal syndromes have primarily been considered at the population and community levels. For example, in Glanville fritillary butterflies, polymorphism in an isomerase gene is such that heterozygotes disperse 70% more often than homozygotes, and because this gene is also associated with differences in clutch size, life span, and other traits, this contributes to colonization-extinction dynamics [2]. An example of community level effects is found in western bluebirds, where the increased aggressiveness of dispersers enables them to out-compete mountain bluebirds in patches they colonize [10].

While such correlations are interesting in the context of population and community dynamics, ecosystems could also be impacted by dispersal syndromes, via resource flux, a measure of ecosystem functioning [11]. In fact, some work has demonstrated that dispersers consume resources differently than non-dispersers; for example, mosquitofish which had dispersed between pools in an experimental stream were four times as efficient at reducing prey abundance after arriving in a new location as are non-dispersers, though this effect attenuated over time [12]. However, this finding was framed in a behavioral context only, ignoring potential ecosystem-level effects. Thus, resource dynamics, and resource consumption in particular, are a potentially unexplored consequence of dispersal syndromes on ecosystems [13].

Detritus consumption by detritivores is a strong determinant of decomposition rate, one of the key fluxes in ecosystems [14,15]. Decomposition of organic matter is especially important in freshwater ecosystems, because it enables terrestrial detritus to subsidize the aquatic food web [16], and shredding of leaf litter by invertebrate detritivores is a key step in the decomposition process [17,18]. Here, we used shredding freshwater detritivores to test whether dispersers differ in their leaf litter consumption rate and thus their contribution to ecosystem function. We used one native and one non-native species of amphipod (Crustacea: Amphipoda), a guild of dominant shredding invertebrates in European streams [19]. Amphipod abundance can drive total terrestrial leaf litter shredding [20,21], however these two species are functionally non-equivalent in their shredding activity [22–24]. After an initial experiment where we allowed individuals to disperse in experimental two-patch landscapes, we examined whether dispersers and non-dispersers (henceforth “residents”) differed in leaf consumption rates.

## Methods

### Study organisms

We used one native amphipods species, *Gammarus fossarum* (Koch), and one non-native amphipod species, *Dikerogammarus villosus* (Sowinsky), in our experiments. *Gammarus fossarum* is very common in headwater streams throughout Switzerland and central Europe [25]. We collected adult *G. fossarum* from the Sagentobelbach stream in Dübendorf, Switzerland (47.39° N, 8.59° E) in November 2016. In the laboratory, amphipods were placed in holding containers of ∼500 individuals and acclimated to 18 °C laboratory conditions for 60 hours, and were provided ad libitum alder (*Alnus glutinosa* (Gaertner)) leaves as food. This was repeated in January 2017 with *D. villosus*, a Ponto-Caspian species which has expanded into central Europe in the last three decades [26], with individuals collected from Lake Constance at Kesswil, Switzerland (47.60° N, 9.32° E). For each species, the experiment was conducted in two steps: a dispersal experiment followed by a leaf consumption experiment. Experimental protocols, including length of dispersal phase and length of consumption experiment, were adapted depending on the species’ activity levels and consumption rates, based on pilot experiments. *Gammarus fossarum* used in the experiment had a mean dry weight of 3.30 mg (s.d. ±1.33), and *D. villosus* had a mean dry weight of 8.59 mg (s.d. ±2.60).

### Dispersal experiment

A common method for examining the causes and consequences of dispersal is to allow organisms to disperse through linked experimental patches ranging from two-patch pairings [27,28] to larger grids or networks [29,30]. The dispersal experiments were run according to the Dispersal Network (DispNet) distributed experiment protocol, detailed in [27]. Briefly, we set up 40 replicates of a two-patch mesocosm system in order to address rates of amphipod dispersal from one to the other patch. The experiment had a factorial design of resource availability (alder leaves vs. no food) and predator cues (fish kairomones vs. no kairmones) in the patch of origin, with each experimental context replicated 10 times. Because we found no effect of the resource or predator cue context on dispersal rates in amphipods [27], we here pooled all data from the different treatments together and only considered the effect of dispersal status (dispersed vs. resident individuals) on subsequent leaf consumption. Residuals from the models (described below) confirmed that no additional variation in leaf consumption rates was explained by experimental context/treatment (Figure S1).

Each patch was a 3 L (198 × 198 mm) polypropelene box, and each pair of patches (one “origin” and one “target” patch, with their relative positions randomized) was connected by 30 cm of silicon tubing with 20 mm diameter. Patches were covered with a black lid to reduce light permeability, while the connection tube was left uncovered; this light difference between patches and matrix rendered the connection tube a hostile matrix, since amphipods are photophobic [31]. Twenty amphipods were placed in each origin patch and allowed to habituate for 30 minutes. We then opened a clamp that had been used to close the connection and amphipods could disperse for a period of 4 ½ hours (*G. fossarum)* or 7 hours (*D. villosus*) before the connection tube was closed again. To confirm that relocation from the origin to target patch was not simply due to routine movement in the course of foraging, but represented dispersal decisions, we also measured movement (gross swimming speed, extracted from videos of the animals using the ‘BEMOVI’ package [32] in R) of residents and dispersers, and found that speed was not correlated with dispersal status (Figure S2).

### Consumption experiment

After the dispersal experiment, amphipods were transferred to new single-patch mesocosms (2 L plastic containers with 0.4 m^2^ of substrate area) to measure leaf litter consumption. The density of amphipods used in the leaf consumption experiment was standardized between dispersers and residents to account for possible effects of density on leaf consumption rates [33]. Thus, from each two-patch system, all dispersers were moved to one new mesocosm, and an identical number of haphazardly-chosen residents was moved to a separate new mesocosm. Densities remained highly correlated at the replicate block level throughout the experiment (*G. fossarum*: r = 0.89, p < 0.001; *D. villosus*, r = 0.53, p = 0.05). Mesocosms were provisioned with 1.5 g (dry weight) of conditioned alder leaves. The leaf consumption experiments were run for 19 (*G. fossarum*) and 12 (*D. villosus*) days, respectively, at which point leaves from the mesocosms were collected and dried for 48 h at 60 °C, then weighed to calculate mass loss from the beginning of the experiment. Amphipods were counted every two to three days throughout the experiments to track mortality; overall, survival was 76.3% for *G. fossarum* and 95.4% for *D. villosus.* These mortality estimates were used to calculate an average daily amphipod density for each mesocosm over the length of the experiment. At the end of the experiment, amphipods were sacrificed and dried for 48 h at 60 °C. The average daily biomass in a mesocosm (mg m^-2^) was then calculated as the average daily density (above) multiplied by the average weight of individuals in the mesocosm. Leaf consumption rates were calculated as the dry weight of leaf litter consumed per milligram of amphipod dry weight per day.

### Analysis

Consumption rates were compared between residents and dispersers of each species separately using linear mixed-effects models with the ‘lme4’ package, version 1.1-18-1 [34], in R version 3.5.0 (R Core Team, Vienna, Austria, 2018). Distributions of consumption rates were positively skewed, so to meet assumptions regarding error structure the *G. fossarum* data were square-root transformed and the *D. villosus* data were inverse-transformed (response = 1/consumption rate) before analysis. For both species, the response was modeled with dispersal status (disperser vs. resident) as a fixed factor, and replicate block (the two-patch experimental metapopulation from which dispersers and residents originated) as a random intercept. The replicate block accounted for all potential differences associated with the experimental metapopulation of origin and density. After building the mixed-effect models, a conditional R^2^ value (accounting for both random and fixed effects) was calculated using the ‘MuMIn’ package, version 1.42.1 [35]. Differences in consumption rates between dispersers and residents were tested using Tukey HSD tests using the ‘multcomp’ package, version 1.4-8 [36].

## Results

For *G. fossarum*, the estimated difference between square-root transformed daily consumption rates of residents and dispersers was 0.020 (standard error of the estimate = 0.121; model R^2^ = 0.38) (Table 1). For *D. villosus*, the estimated difference between inverse-transformed daily consumption rates of residents and dispersers was 0.208 (standard error of the estimate = 0.063; model R^2^ = 0.82), which was significant according to post-hoc testing (z = 3.31, p < 0.001, Table 1). Dispersing *D. villosus* had similar biomass-adjusted consumption rates to *G. fossarum*, and approximately three times higher than non-dispersing *D. villosus* (Figure 1).

**Table 1.** *(next page)* Results from the linear mixed-effects models of biomass-adjusted consumption rates as a function of dispersal status, for *Gammarus fossarum* (n=73 mesocosms) and *Dikerogammarus villosus* (n=53). Estimates and their standard errors are drawn from linear mixed-effects models, and z-and p-values for the effect of dispersal status are from Tukey’s HSD tests; variance associated with the random factor of replicate blocks, and its standard deviation, is reported in italics.

|  | Coefficient | Std. Error/<br>Std. Dev. | z | p |
| --- | --- | --- | --- | --- |
| ***G. fossarum* (square-root transformed daily consumption)** |  |  |  |  |
| Intercept (residents) | 1.951 | 0.110 |  |  |
| Dispersers | 0.019 | 0.121 | 0.163 | 0.87 |
| *Variance due to replicates:* | *0.156* | *0.395* |  |  |
| ***D. villosus* (inverse-transformed daily consumption)** |  |  |  |  |
| Intercept (residents) | 0.666 | 0.080 |  |  |
| Dispersers | 0.209 | 0.063 | 3.311 | < 0.001 |
| *Variance due to replicates:* | *0.125* | *0.354* |  |  |

**Figure 1.**
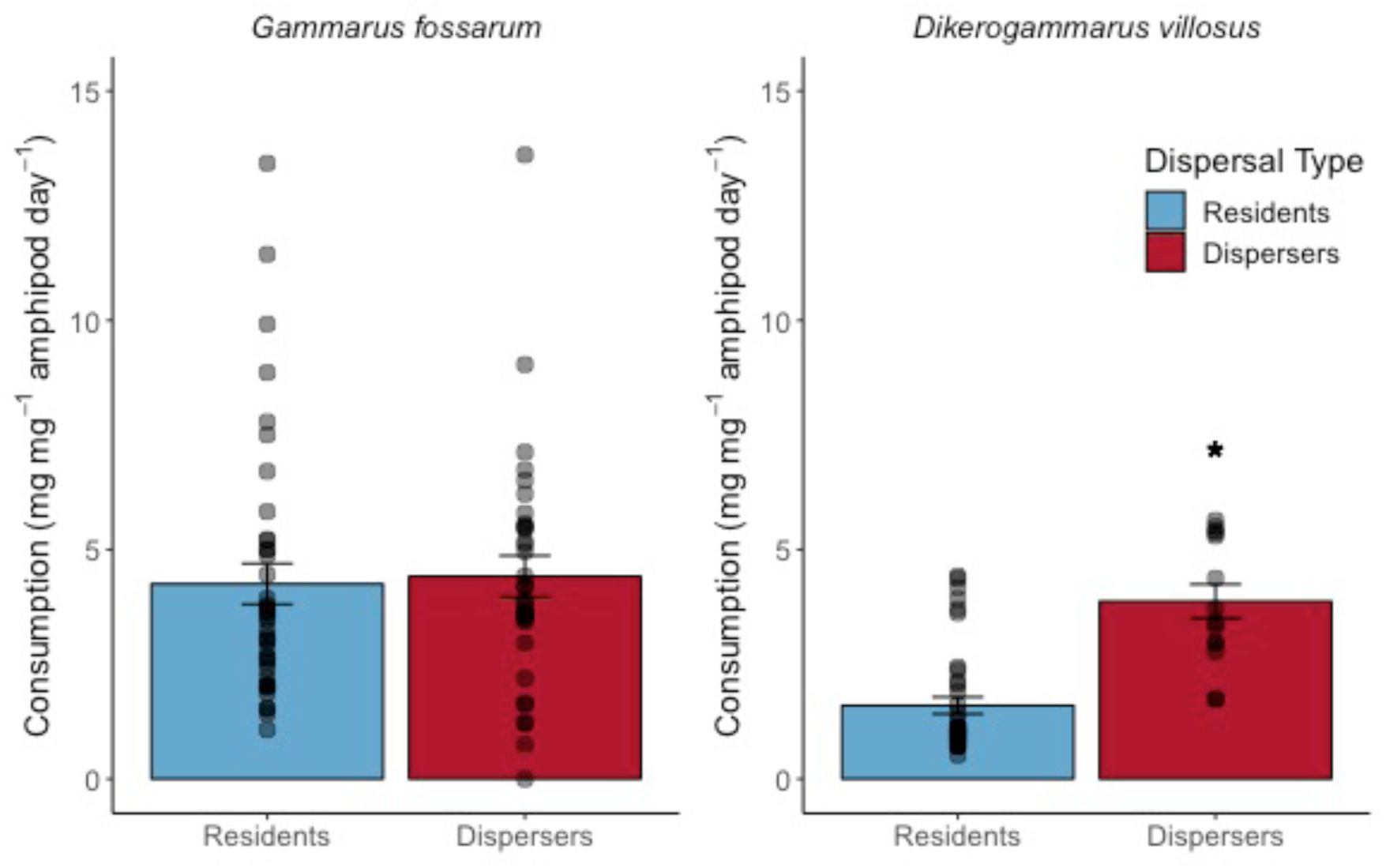
Daily average leaf litter consumption by dispersing and non-dispersing (“resident”) amphipods of *G. fossarum* (n = 73 mesocosms) and *D. villosus* (n = 53), adjusted for biomass of the individuals in each experimental replicate. Error bars show standard error of the mean, and gray dots show raw data points from experimental mesocosms. Asterisk shows a significant difference (p < 0.05) between consumption rates of dispersers and residents according to a linear mixed effect model.

## Discussion

We identified a dispersal syndrome with consequences for ecosystem functioning in a non-native but not a native amphipod species: *D. villosus* dispersers consumed leaf litter at roughly three times the rate of residents, while there was no difference in leaf consumption rate with dispersal status in *G. fossarum*. To date, most research addressing consumption rates in relation to dispersal status or range fronts has been in a behavioral context, addressing personality and aggression as contributions to predator-prey interactions [12,37,38], for example. To our knowledge there has been little research into consumption of basal resources as a component of nonrandom dispersal. This is despite the importance of such traits to energy flows through food webs and ecosystems. Furthermore, differences in traits that may depend on resource consumption – such as size, metabolism, and growth rates [2,8] – with dispersal propensity render resource consumption a logical component of a dispersal phenotype, and thus one which could have consequences for energy fluxes through food webs and ecosystems.

Our study species are omnivorous aquatic invertebrates, which despite a wide diet breadth contribute the bulk of leaf litter processing in central European headwater streams [20]. Our results show that in *D. villosus*, dispersers make a greater contribution to the detritus-based pathway integrating terrestrial energy into the food web than do residents. This species also has lower overall contributions to leaf litter processing than *G. fossarum* [22–24], but we suggest that both species identity and dispersal status of individuals within a species could jointly determine their contribution to ecosystem function.

Predicting these populations’ contributions to ecosystem function is important because *D. villosus* has been deemed one of the 100 worst invaders in European freshwater ecosystems [39]. Because the non-native species is currently undergoing a range expansion, the signature of either trade-offs for increased dispersal ability or selection for success in new habitats is likely more prominent than in populations which are in their range core (such as the *G. fossarum* populations used in our experiment), consistent with spatial selection theory [40]. Identifying whether this is true or whether the dispersal syndrome is consistent across the range of *D. villosus* would require performing experiments with *D. villosus* from its range core in the Ponto-Caspian region. This would also address whether it is appropriate to make interspecific comparisons of this and other phenotypic traits using populations with different recent dispersal/range expansion histories, depending on the research question.

Regardless, how non-native species will affect ecosystem function is a central question in the era of global change and increased connectivity [41]. As the distribution of suitable habitat is altered and human activity continues to contribute to global organismal dispersal, the potential effects of phenotype-dependent dispersal should be considered when attempting to predict impacts on ecosystem function. This may be challenging, because it means that predictions made based on species contributions to ecosystem function in their range core may not be valid at the edges of their range expansions [41]. However, considering the increasing evidence of how dispersal phenotypes can alter system dynamics, it is crucial to extend this understanding into the realm of ecosystem function.

## Acknowledgements

The authors thank Samuel Hürlemann, Remo Wüthrich, Georg Flückiger, and Sascha Brunner for help in the lab and field, and thank Felix Moerman for comments on an early version of the manuscript. We thank the members of DispNet for the collaboration that led to this experiment. This is publication ISEM-YYYY-XXX of the Institut des Sciences de l’Evolution -- Montpellier.

## Author contributions

EAF conceived the dispersal experiment and all authors together designed the consumption experiment. CJL and EAF ran the experiments. CJL analyzed the data and drafted the manuscript. All authors contributed to revisions, gave final approval for publication, and agreed to be held accountable for the work within the article.

## Data accessibility

Data will be made available on Dryad and code will be posted to GitHub upon the manuscript’s acceptance; contact the authors (information on cover page) in the meantime to request access to data and code.

## Ethical statement

No ethics approval was required for this experiment. Work with non-native species was carried out according to the laws of Switzerland.

## Supplementary Material

**Figure S1.**
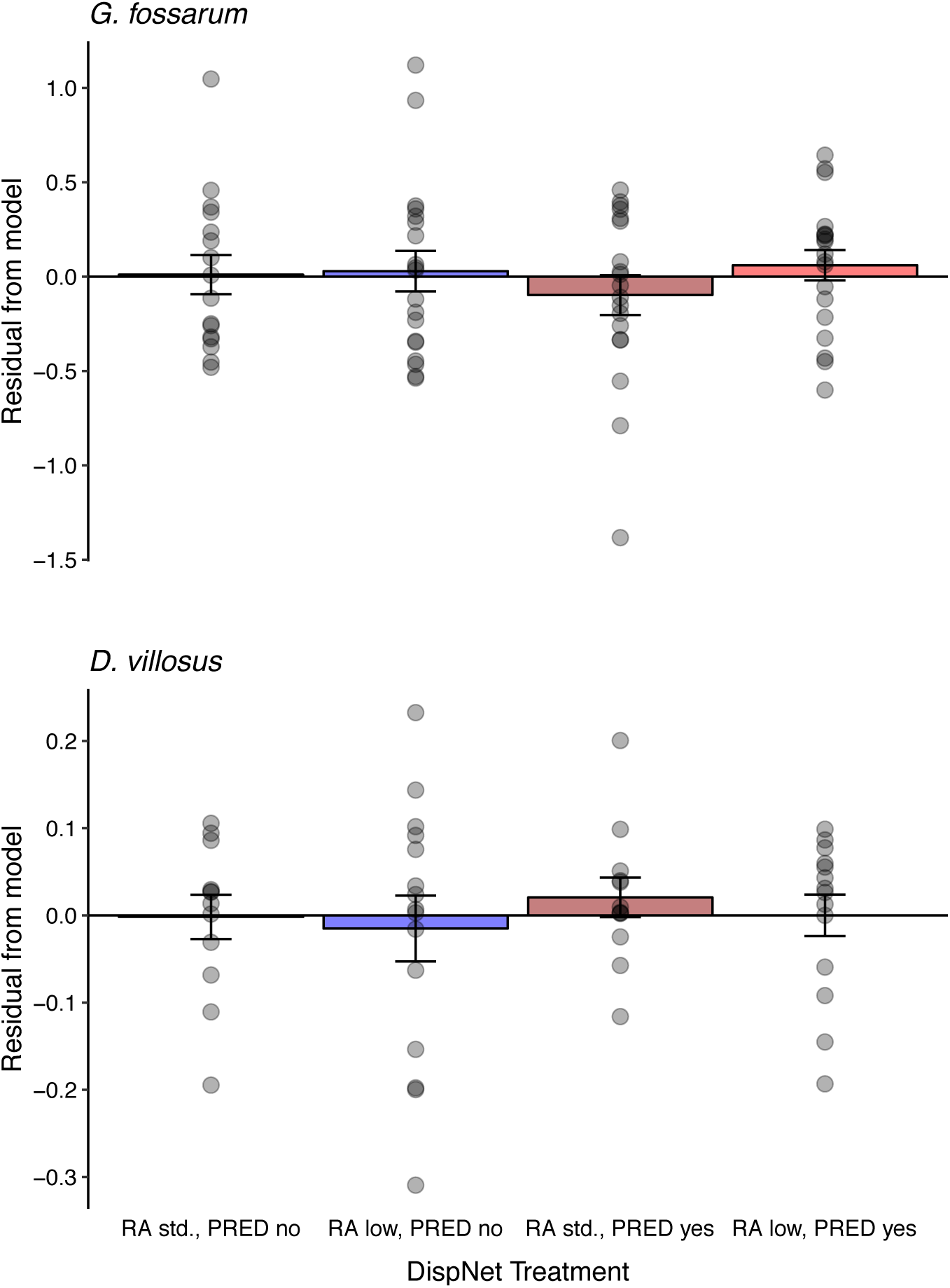
Model residuals from the mixed-effect models (transformed consumption rate ∼ dispersal status + (1|replicate block)) plotted against treatments from the dispersal experiment: RA = resource availability (standard or low), PRED = predator cues (no or yes). Linear models of residuals as a response of dispersal experiment treatment showed no significant effects (*G. fossarum*: F_3,69_ = 0.49, p = 0.69; *D. villosus*: F_3,49_ = 0.25, p = 0.86). Error bars show standard error of the mean, and gray points show residuals from individual experimental replicates.

**Figure S2.**
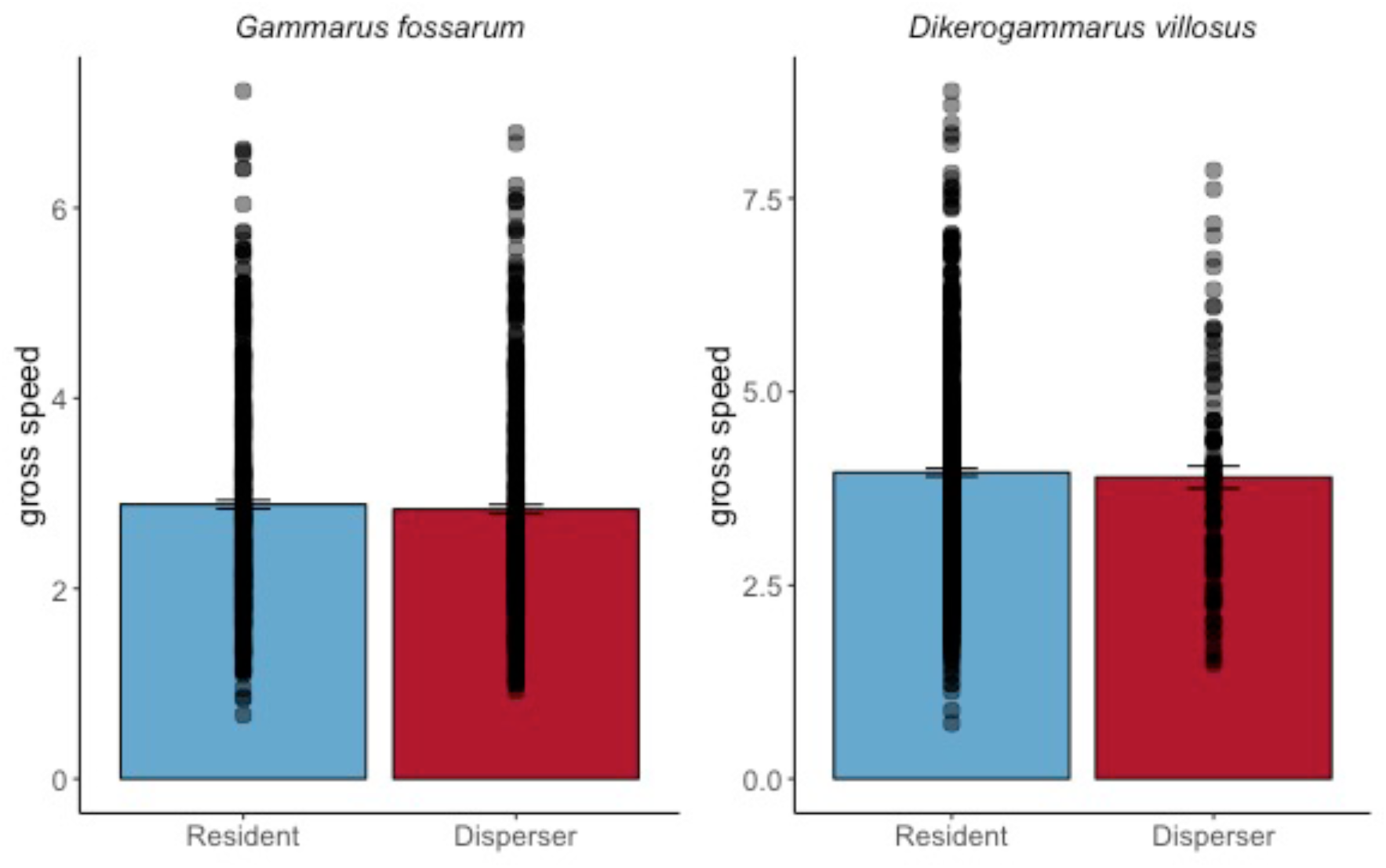
Gross swimming speed of residents and dispersers, from video analysis using the ‘BEMOVI’ package in R. Before being placed into the consumption mesocosms, residents and dispersers were (separately) placed into an experimental arena and allowed to move freely for three minutes. Each time an amphipod moved it was detected it was given an object identifier and the movement was described; gray dots in the figure represent each movement, and error bars show the standard error of speed for residents and dispersers. There were no significant differences in swimming speed between residents and dispersers based on simple linear models in either *G. fossarum* (F_1,1109_ = 0.57, p = 0.44) or *D. villosus* (F_1,824_ = 0.17, p = 0.68).

